# Effect of flicker-induced retinal stimulation of mice revealed by full-field electroretinography

**DOI:** 10.1101/2024.12.27.630543

**Authors:** Milan Rai, Yamunadevi Lakshmanan, Kai Yip Choi, Henry Ho-lung Chan

## Abstract

**Purpose:** To investigate the effects of brief flickering light stimulation (FLS) on retinal electrophysiology and its blood flow in normal C57BL6J mice.

**Methods:** Retinal blood flow (RBF) and full-field electroretinography (ffERG) were measured before and after a 60-second long FLS (12 Hz, 0.1 cd·s/m^2^) in a cohort of 8-12 weeks old C57BL6J mice (n=10) under anaesthetic and light-adapted conditions. A separate set of age-matched mice (n=9) underwent RBF and ffERG measurements before and after steady light stimulation (SLS) at 1 cd/m^2^ under similar conditions. The changes in RBF (arterial and venous flow) and ffERG responses (amplitudes and implicit times of a- and b-wave) were analyzed.

**Results:** FLS significantly increased both arterial (p=0.003) and venous (p=0.018) blood flow as well as b-wave amplitudes (p=0.017) compared to SLS, which did not have any significant changes in both RBF and ERG. However, no significant differences were found in other ffERG responses (amplitude and implicit time of a-wave and b-wave implicit time) between the two groups after light stimulation. An increase in b-wave amplitude was positively associated with increase in both arterial (r=0.655, p=0.040) and venous blood flow (r=0.638, p=0.047) in the FLS group.

**Conclusions:** Transient FLS induced a significant increase in both RBF and electro-retinal activity, but such increase was not observed after SLS. Our results suggest the role of FLS, which exerts metabolic stress on the retina, in triggering retinal neurovascular coupling.

## Introduction

The robust vascular system intrinsically maintains an adequate blood supply to neural tissues, which is essential for meeting the high metabolic demands associated with neural activities [1-4]. The retina, like other structures of the central nervous system, is a complex multi-layered tissue which consists of different types of neurons, including photoreceptors, bipolar cells, and retinal ganglion cells, with high metabolic needs. These retinal neurons require a large amount of oxygen and glucose supplied by the choroidal and retinal vasculature. It implies that increments in the metabolic demands of these neurons have to be supported by an augmented supply of oxygenated blood and nutrients. Retinal stimulation with flickering light has been reported to substantially increase the retinal metabolism [5]. Both preclinical and clinical studies have reported substantial retinal vasodilation and increased retinal blood flow (RBF) in response to high retinal metabolic activity induced by flickering light stimulation, as measured by techniques such as laser speckle flowgraphy, dual beam bi-directional Doppler Fourier-domain optical coherence tomography (OCT), and retinal vessel analyser [6-18]. However, the retinal vascular caliber changes and increased RBF do not directly relate to the physiological status of retinal cells, as there is no direct reflection of the actual variations in retinal functional status in response to the FLS. Hence, there is limited information regarding the corresponding changes in the cellular activity (retinal function) elicited by FLS. Furthermore, the relationship between retinal vascular caliber changes and the physiological status of retinal cells remains unclear.

We proposed that retinal activity would be increased after retinal stimulation by flickering light and this can be readily assessed using electroretinogram, a tool that allows measuring electrical activity from distinct retinal layers. Therefore, this study reports the effect of FLS on retinal electrophysiological activity along with the RBF through application of full-field electroretinogram (ffERG) and Doppler spectral-domain optical coherence tomography (SD-OCT) in normal adult mice.

## Materials and Methods

### Animals

Nineteen C57BL/6J mice (age: 8-12 weeks, 10 males and 9 females), obtained from the Centralised Animal Facility of The Hong Kong Polytechnic University, were housed in a temperature-controlled room (21-22^°^C) under normal lighting conditions (approximately 200 lux) with a 12/12-hour light/dark cycle. Mice were given ad libitum access to food (Pico Lab diet 20 (5053); PMI Nutrition International, Richmond, IN, USA) and water. All experimental procedures were carried out in adherence with the ARVO Statement for the Use of Animals in Ophthalmic and Vision Research. The study protocol was approved by the Animal Ethics Sub-committee of The Hong Kong Polytechnic University (Animal Subjects Ethics Sub-committee approval number: 18-19/58-SO-R-OTHERS).

### Measurement of retinal blood flow

RBF was evaluated using an annular Doppler SD-OCT (Envisu R2210; Bioptigen, Morrisville, NC, US) as described previously [19,20]. This OCT utilized the Doppler shift phenomenon to enable the direct visualization of flow towards and away from the objective. At first, the circular scanning with a diameter of 0.5 mm was positioned with its center located at the center of the optic nerve head. Doppler shifts appeared as red and blue signals, representing arterial (towards the objective) and venous (away from the objective) blood flow, respectively. Subsequently, ten OCT B-scan images, featuring both red and blue signals, were captured. The regions displaying Doppler shifts were then cropped using FIJI software (https://imagej.net/software/fiji/) [19,20] for analysis. The saturation of each cropped B-scan image was adjusted using the color threshold algorithm (available in same software under the ‘Adjust’ sub-menu of the ‘Image’ category) to include all red and blue signals in the retinal blood vessels. After removing the noise present in regions other than the blood vessels, both red and blue signals were separately quantified in terms of pixels using ‘Measure’ function of the ‘Analyze’ category of the software. The values from ten B-scans were then averaged to determine the final red/blue pixels. The mean pixel count serves as a surrogate measure of retinal arterial (red pixels) and venous (blue pixels) calibers (per se the blood flow), with a higher pixel count indicative of greater RBF, and a lower count indicating less RBF.

### Full-field Electroretinography

The ffERG was recorded using a Ganzfeld ERG system (Q450; RETI Animal, Roland Consult, Brandenburg an der Havel, Germany). The animal, after pupil dilation, was placed on a heating pad to maintain the body temperature at around 37^°^C. A drop of lubricating gel was applied on the cornea to prevent corneal dehydration. For ffERG recording, a 2-mm-diameter gold ring electrode (Roland Consult) was placed on the cornea of each eye as an active electrode. Needle electrodes (Item No. U51-426; GVB-geliMED, Bad Segeberg, Germany) were inserted into lateral canthus of each eye and into the upper base of the tail as reference and ground electrodes, respectively. An impedance of less than 5 kΩ was maintained for all electrodes throughout the recording period. The ERG responses were elicited by presenting a brief white LED flash (each flash duration: 4 ms) of 3.0 cd·s/m^2^ under the Ganzfeld stimulator. The signals were amplified and filtered with a bandpass of 0.1 -1000 Hz. Twenty-five ERG responses (interstimulus intervals of 1 sec) were recorded and averaged. The amplitudes and implicit times of a-wave and b-wave were extracted for analysis.

### Flickering light stimulation

A custom-built device with a LED source was used to produce white flickering light stimulation (FLS). Square-wave flickering light (frequency:12 Hz, intensity: 0.1 cd·s/m^2^) was selected for retinal stimulation, as flickering light at this frequency and low intensity has been reported to effectively increase RBF in rod-dominant retina [16,21]. As Werkmeister et al. reported a strong increase in RBF during and after 60-second of 12 Hz FLS, the current study employed the same stimulation frequency and duration [21].

### Procedures

#### Experiment 1 – Flickering light effect on retinal blood flow

Ten mice were used to investigate the effect of FLS on RBF. Following overnight dark adaptation, animals were then anesthetized by intraperitoneal injection of a cocktail containing 90 mg/kg ketamine (Alfasan International B.V., Woerden, Holland) and 12 mg/kg xylazine (Alfasan International B.V.). Animal preparations and procedures were performed under dim red illumination. Briefly, a drop of topical anesthetics (Provain-POS 0.5% wt/vol eye drops; URSAPHARM, Saarbrücken, Germany) was applied to the eyes and the pupils were dilated by using a mydriatic agent (Mydriacyl 1% eye drops; Alcon-Couvreue, Puurs, Belgium). One randomly selected eye was light adapted for 10 minutes (1 cd/m^2^), using the custom-built device and then baseline doppler SD-OCT was taken as described above. Subsequently, retina was stimulated using a 12 Hz flickering light of 0.1 cd.s/m^2^ for 60 seconds. Promptly after the cessation of FLS, doppler OCT was repeated. The background luminance of 1 cd/m^2^ was maintained throughout the experiment except during FLS. Figure 1 shows the experimental procedures of Experiment 1.

**Fig 1.**
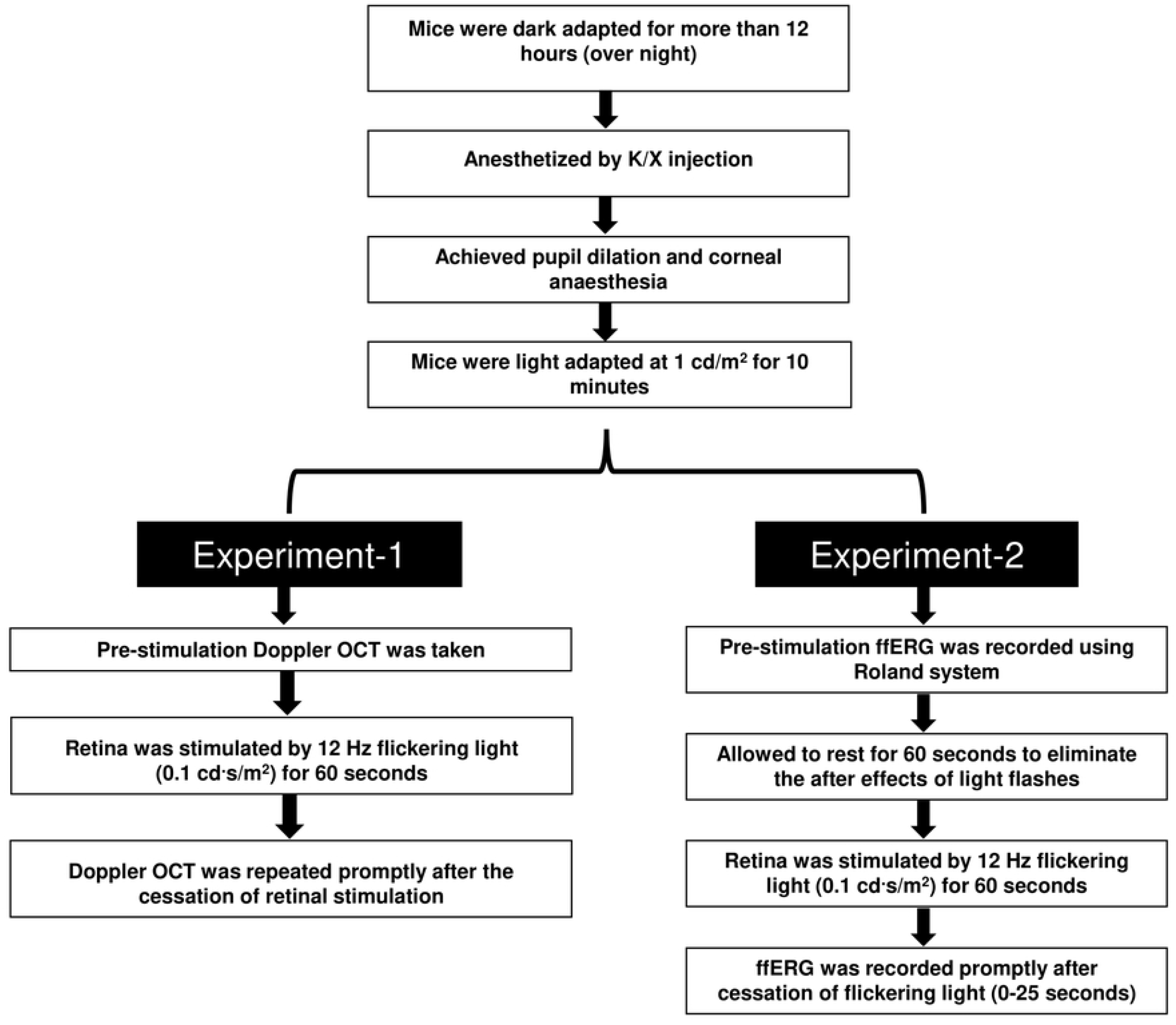
Flow diagram showing the experimental procedures of Experiments 1 and 2. Doppler OCT and FfERG recordings of light-adapted anesthetized C57BL6J mice were performed before and promptly after the cessation of 60 sec long flickering light stimulation (12 Hz, 0.1 cd·s/m^2^).

#### Experiment 2– Flickering light effect on retinal electrophysiological responses

One week following Experiment 1, same cohort of ten mice were used to measure the FLS induced change in electro-retinal activity. Using the same anesthetic and mydriatic protocols, overnight dark-adapted mice were firstly light-adapted at 1 cd/m^2^ light for 10 minutes. A baseline ffERG recording was taken, and animals were then given a 60-second resting period to eliminate the aftereffects of ERG light flashes. Subsequently, the retina was exposed to the FLS for 60 seconds, and ffERG recordings were repeated promptly (within 25 seconds) after the cessation of FLS. The background luminance was constantly maintained at 1 cd/m^2^ throughout the experiment, with the exception of 60-second FLS period. Figure 1 depicts the experimental procedures of Experiment 2.

#### Experiment 3 – Steady light effect on retinal blood flow and retinal electrophysiological responses

In order to assess whether the increase in RBF and ffERG responses were specifically caused by FLS or by random variation, a control group was used. RBF and ffERG responses were obtained in a new cohort of nine mice following experimental protocols similar to Experiments 1 and 2 respectively. However, in this experiment, mice were continuously exposed to the steady light of 1 cd/m^2^ (approximately equivalent to the light energy produced by FLS of 0.1 cd.s/m^2^) for 60 seconds. The ffERG recording was carried out one week after RBF measurement to allow sufficient rest for mice. Both procedures were conducted under red-dim illuminations. A Supplementary appendix (S1 Appendix) illustrates the calculations converting the luminous intensity of the flickering light to equivalent steady-state value.

## Statistical Analysis

The data from all experiments had no outliers and followed a normal distribution (Shapiro-Wilk test, p > 0.05). Data are presented as mean ± standard error of the mean (SEM). Mixed-model ANOVA was applied to assess the difference in RBF (arterial and venous blood flow) and ffERG responses (amplitudes and implicit times) between two groups and also within the groups measured before and after the flickering or steady light stimulation with Bonferroni’s pairwise post hoc comparisons. The relative change in RBF (ABF and VBF) and ffERG responses (amplitudes and implicit times) was calculated as percentage change from baseline. Then, unpaired two-tailed t-test was applied to compare the difference between FLS and SLS groups. Pearson’s correlation test was employed to determine the relationship between percentage change in ffERG responses with percentage change in ABF and VBF. All statistical analyses were conducted by using SPSS 29.0 (IBM Corp., Armonk, NY, USA). A p-value < 0.05 was considered statistically significant.

## Results

Figs 2A and 2B show the representative doppler B-scans (imaged before and after the light stimulations) related to RBF of FLS and SLS groups, respectively. The mean values (in terms of number of pixels) of ABF (Fig. 2C, F) and VBF (Fig. 2D, G) and their corresponding percentage change (Fig. 2E, H) are also presented in fig 2.

**Fig 2.**
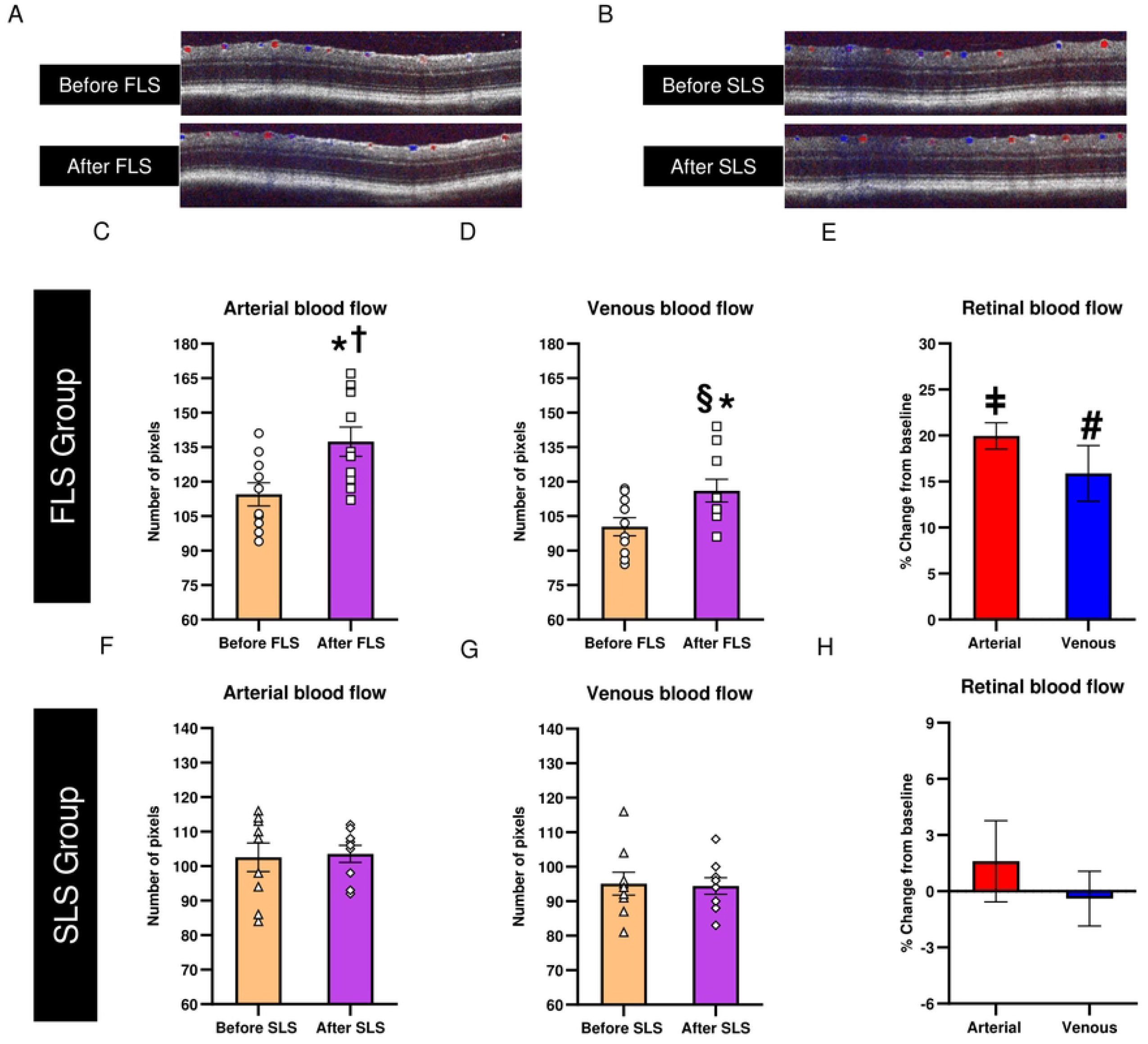
Effects of FLS and SLS on the RBF, assessed by SD-OCT Doppler system. (A, B) Doppler B-scans of one representative mouse retina, from FLS and SLS group measured before and after stimulation. Mean values of (C, F) ABF and (D, G) VBF assessed before and after stimulation from each group and mean (E, H) percentage changes of ABF and VBF from corresponding baseline from each group. Error bars represent standard error of mean (SEM). Data points represent individual mouse data. ^*^ P < 0.001 when compared with relative baseline, ^†^p < 0.001 when compared with After SLS (arterial) and, ^§^ P = 0.001 when compared with After SLS (venous) using mixed-model ANOVA with Bonferroni post hoc correction, ^‡^p < 0.001 and ^#^ P < 0.001 when compared with percentage change in ABF and VBF, respectively in SLS group using unpaired two-tailed t-tests.

For both ABF and VBF, significant differences between FLS and SLS group were detected [Mixed-model ANOVA: (ABF: time effect: P < 0.001, between groups: P = 0.003 and interaction effect: P < 0.001) and (VBF: time effect: P = 0.001, between groups: P = 0.018 and interaction effect: P < 0.001)]. Furthermore, RBF after FLS, was significantly greater as compared with RBF after SLS (Mixed-model ANOVA with Bonferroni post hoc test: P < 0.001 for ABF and P = 0.001 for VBF) and also, with pre-flickering baseline measurements (both P < 0.001). Similarly, significant differences were detected in the percentage changes in ABF and VBF between the FLS and SLS groups (Unpaired t-test: p < 0.001 for both ABF and VBF). However, SLS showed no significant changes in both ABF or VBF when compared with pre-flickering baseline measurements (P = 0.642 for ABF and P = 0.796 for VBF).

Figs 3A and 3B illustrate the traces of averaged ffERG responses recorded at two time points (before and after stimulation) from one representative animal from FLS and SLS group, respectively. The averaged amplitudes (Fig. 3C, E, G, I) and implicit times (Fig. 3D, F, H, J) of both a- and b-waves, measured before and after stimulation, and the corresponding percentage change (Fig. 3K, L, M, N) from baseline for both FLS and SLS group are also presented.

**Fig 3.**
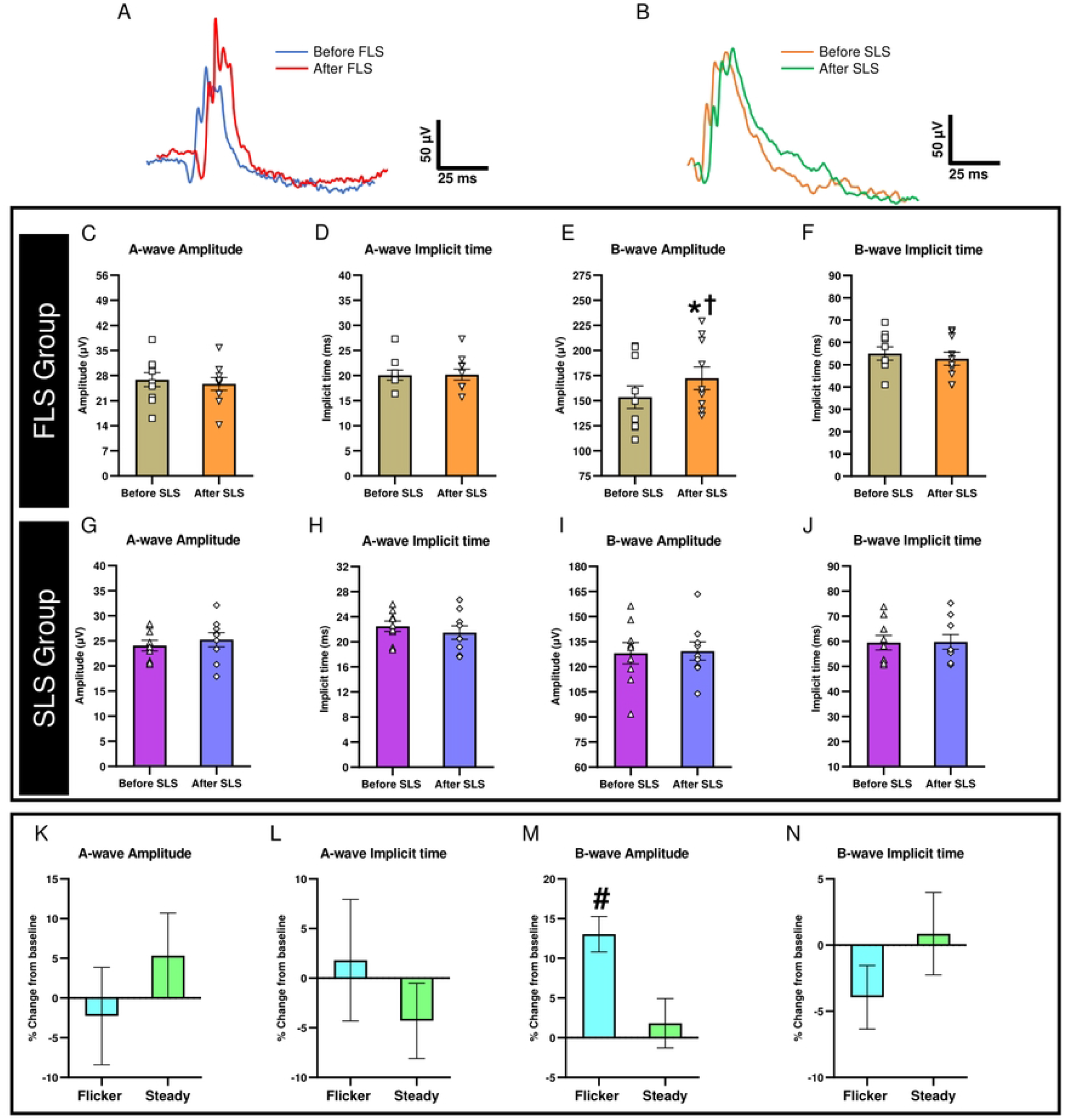
Effects of FLS and SLS on ERG responses. The averaged traces of ERG responses of one representative mouse from (A) FLS group and (B) SLS group measured before and after stimulation. Mean values of ERG responses (C, G) a-wave amplitudes, (D, H) a-wave implicit times, (E, I) b-wave amplitudes, (F, J) b-wave implicit times and (K, L, M, N) the corresponding percentage changes from baseline. Each data point represent response from an individual mouse. Error bars represent SEM. ^*^ P < 0.001 when compared with baseline and ^†^ P = 0.004 when compared with b-wave amplitude measured after SLS using mixed-model ANOVA with Bonferroni post hoc correction and ^#^ P = 0.010 when compared with SLS group (percentage change in b-wave amplitudes from baseline) using unpaired two-tailed t-test.

The amplitudes of b-wave showed a significant difference between FLS and SLS group (Mixed-model ANOVA: time effect: P < 0.001; between groups: P = 0.017; interaction effect: P < 0.001). However, there were no significant differences in other ERG parameters between two groups [Mixed model ANOVA: implicit times of the b-wave (time effect: P = 0.389; between groups: P = 0.168; interaction effect: P = 0.287), amplitudes (time effect: P = 0.991; between groups: P = 0.424; interaction effect: P = 0.348) and implicit times of a-wave (time effect: P = 0.567; between groups: P = 0.133; interaction effect: P = 0.495)]. The b-wave amplitudes measured after FLS was significantly greater compared to the amplitudes of b-wave recorded after SLS (post FLS vs post SLS, Mixed-model ANOVA with Bonferroni post hoc test: P = 0.004). In contrast, other ERG parameters measured after FLS were not different from those measured after SLS [Mixed-model ANOVA with Bonferroni post hoc test: b-wave implicit times (P = 0.107), a-wave amplitudes (P = 0.853) and a-wave implicit times (P = 0.403)]. The percentage change in b-wave amplitudes was also significantly increased in FLS as compared with SLS group (Unpaired t-test: P = 0.010). However, no significant differences were found in percentage change in b-wave implicit times (Unpaired t-test: P = 0.241), a-wave amplitudes (Unpaired t-test: P = 0.362), and a-wave implicit times (Unpaired t-test: P = 0.410) between two groups. FLS significantly increased the b-wave amplitudes as compared with its pre-flickering baseline (P < 0.001), and no significant change was observed in SLS group (P = 0.668). In addition to this, no significant differences were detected in other ERG responses [ Mixed-model ANOVA with Bonferroni post hoc test: b-wave implicit times (P = 0.168 for FLS and P = 0.883 for SLS), a-wave amplitudes (P = 0.487 for FLS and P = 0.520 for SLS) and a-wave implicit times (P = 0.935 for FLS and P = 0.390 for SLS)] compared to corresponding baseline in both groups.

Notably, a significant positive correlation was found between the percentage change in b-wave amplitudes and percentage change in ABF (r = 0.655, P = 0.040) and VBF (r = 0.638, P=0.047) in FLS group (Figure 4) but not in SLS group (ABF: r = -0.042, P = 0.914; VBF: r = -0.132, P = 0.735).

**Fig 4.**
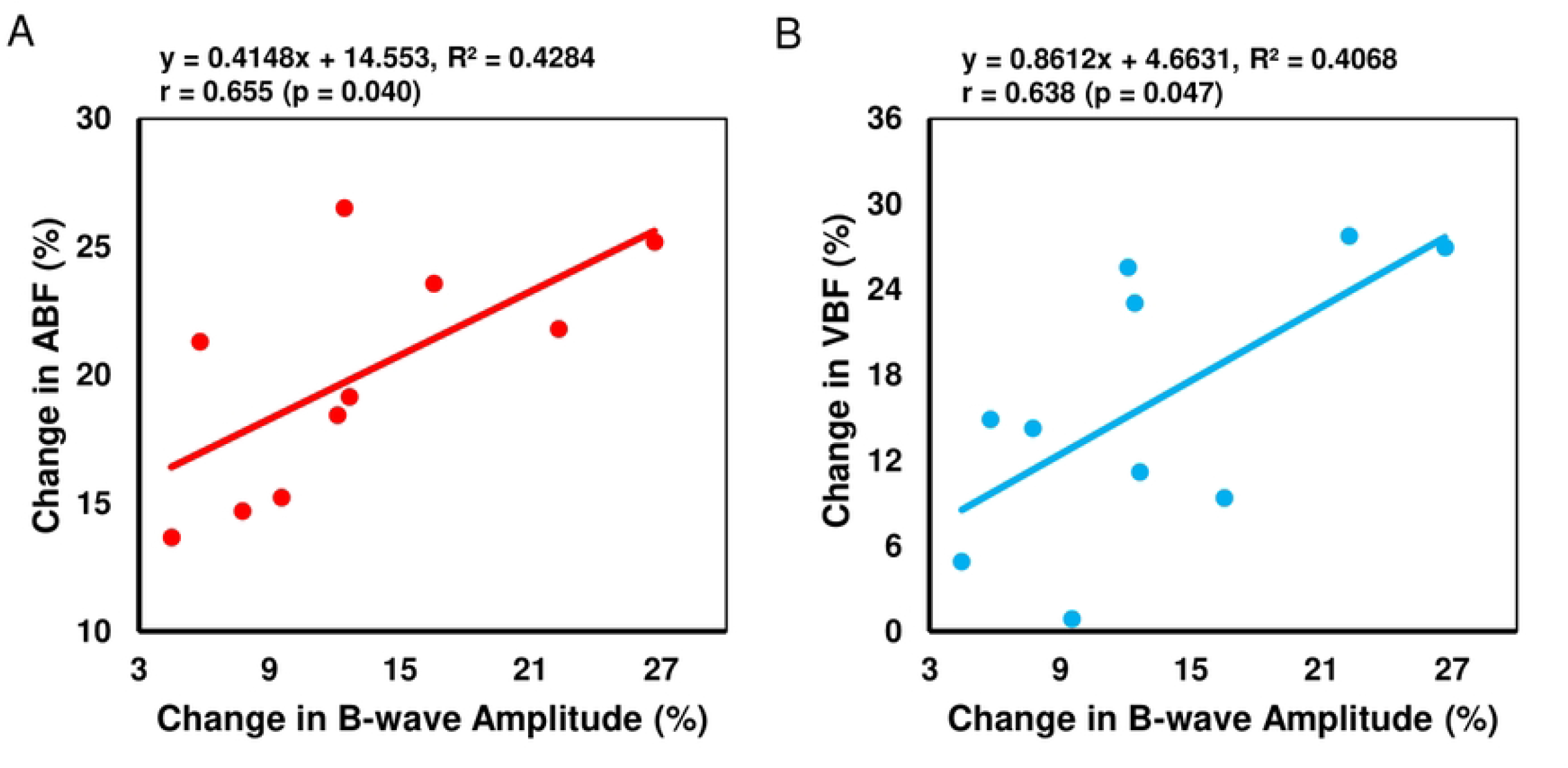
Correlations of percentage change in b-wave amplitude with (A) percentage change in ABF and (B) percentage change in VBF.

## Discussion

To our knowledge, the present study is the first to report the effect of FLS on electrical activities together with blood flow change in mice retina. The findings suggested that the FLS increased the electrophysiological responses, originating primarily from the mid-retinal layer (i.e., bipolar cells, mainly represented by b-waves) and the RBF. Previous studies largely reported the effects of flickering light on retinal blood flow velocity and/or blood volume [6-18]. However, no study has yet reported the changes in retinal electrophysiological responses promptly after flickering light stimulation. Therefore, the current study employed ERG, in addition to RBF measurement, to investigate the flicker-induced neuronal activities in the retina, thus reporting the changes in both vascular and neuronal components.

In this study, the average increase in ABF and VBF were 19.96±1.41% and 15.89±3.02 % respectively. A previous study using C57BL6J mice reported a maximum increase of 32.5 ± 5.0% in RBF during 3 minutes of retinal stimulation (12 Hz, 30 lux) [16]. Another study detected an increase of 22.79-26.37% in relative RBF during 20 seconds of FLS (10 Hz, 1200 lux) using similar mouse strain [22]. It is apparent that the increase in RBF varied between studies and this could be attributed to the differences in measurements used (laser speckle flowgraphy vs Doppler OCT), stimulus parameters (in terms of duration, frequency, luminance/intensity of flickering light) and methodology (during vs after stimulation) employed. Our results showed that the b-wave amplitude measured promptly after FLS was significantly increased by 13.04±2.23%, although post-flickering b-wave implicit time did not differ significantly from the corresponding baseline value. On exposure to SLS using similar light energy-matched intensity, the post-flickering b-wave amplitudes were similar to the corresponding baseline. These suggested that the FLS (in terms of its temporal characteristics) induced significant effects on retina by increasing the RBF and ERG response as compared with SLS. As FLS has been hypothesized to induce neuro-associated vascular changes in the retina, an increase in flicker induced RBF lead to an alteration in retinal electro-activity and this corresponds with an increase in b-wave amplitudes as observed in the present study.

The neuro-associated vascular changes observed in this study may be related to the neurovascular coupling, which is an auto-regulatory mechanism linking the transient increase in metabolic demands of local neural tissues with subsequent augmented blood supply to those tissues [1]. Previous studies have reported that the retinal stimulation by flickering light induces neurovascular coupling-related activities whereby retinal vasodilation occurs, increasing the retinal blood supply to sustain the high metabolic demands of retinal cells caused by the repeated light stimulation [6-18]. The augmented blood supply nourishes the retinal cells by providing higher amounts of nutrients and electrolytes and, hence, enhancing the physiological responses of retinal neurons, which is believed to be reflected by the electro-retinal activities measured by ERG in response to FLS.

Our results suggested that the increment in post-flickering b-wave amplitude was accompanied by a flicker-induced increase in RBF. Furthermore, previous studies reported a rapid loss of ERG in hypoxemic [23], hypoglycemic [24], and ischemic conditions [25], indicating the critical role of adequate blood supply in sustaining normal retinal physiology. However, our results did not demonstrate a significant change in ffERG a-wave after FLS, suggesting that the physiological mechanism of flicker-induced augmented blood flow to the retina is mainly related to the increased inner retinal metabolism, as inner retinal neurons undergo net depolarization and increased firing rate during flickering light exposure [5]. Buerk et al. reported an increase of K^+^ production correlated with the blood flow changes at the surface of the optic nerve head in response to FLS [26]. Dmitriev et al. recently reported a sustained increase in K^+^ concentration in the inner retina, when an isolated mouse retina was stimulated with flickering light, which suggested increased metabolic demands in that region [27]. The continuous transport of K^+^ requires an additional supply of adenosine triphosphate (ATP) molecules to function against the concentration gradient. An increase of RBF is believed to meet the augmented metabolic demand caused by high demand of ATP, which may, in turn increase the physiological activity of the retinal cells. Supporting this, Kornfield and co-workers reported the maximum increase in RBF within the intermediate and deep vascular layers, which supply the terminals of bipolar cells [10]. Notably, our study demonstrated that FLS on the retina led to a significant increase in electrophysiological responses, predominantly from the mid-retinal region, which is likely the site of the retinal bipolar cells, thereby supporting the hypothesis of a metabolic and functional link between RBF and electro-retinal activity.

One of the limitations of this study is that ketamine-xylazine cocktail was used as a general anesthetic, which has been reported to affect the hemodynamic conditions by lowering systemic blood pressure in mice. Nonetheless, even under the influence of ketamine-xylazine, Tamplin et al., using the same mouse strain (C57/Bl6J) as the present study, also reported a significant increase in RBF compared to baseline during flickering light exposure [22]. Furthermore, as both baseline and post-flickering ERG responses were recorded from the same animals under identical conditions, the impact of physiological variations due to general anesthesia should be minimal. Another limitation of this study includes the distinct application of low-intensity flicker setting (Experiment 1 and 2) and low-luminance steady-state condition (Experiment 3). The intended luminance level for the SLS was 1.2 cd/m^2^ to match the luminous energy produced by 0.1 cd·s/m^2^. However, due to technical limitation of the employed device, a luminance level of 1.0 cd/m^2^ was utilized. However, this distinction, despite precluding direct comparability between two values, entails that both the flickering and steady-state conditions utilized low-level stimuli. This could have had a very mild impact on the findings of our study.

By applying our experimental protocols (both RBF and ERG), further studies can be carried out to explore neuro-associated vascular changes that mimic ocular conditions related to retinal vasculature, such as hypertensive and diabetic retinopathies. Comprehending the relationship between retinal neural and vascular component through applications of ERG and RBF, respectively, could facilitate a deeper understanding on how the neural activity and vascular responses are coupled under the influence of flickering light stimuli. And, this may initiate potential applications in clinical practice.

In conclusion, this study quantified the changes in electro-retinal responses in response to flickering light stimulation and demonstrated an association between flicker-induced changes in RBF and ERG responses in wild type normal mice. The increased retinal electrophysiological response from the mid-retinal layer following a brief flickering light stimulation is positively correlated with the increase of retinal blood flow, which is possibly related to the effect of neurovascular coupling.

## Acknowledgments

The authors thank Maureen Boost for proofreading the manuscript.

## Supporting information

**S1 Appendix. Appendix file showing the conversion of luminous intensity of the flickering light to equivalent steady-state value**.

**S2-Retinal blood flow data sets. Excel file containing all data underlying the findings related to retinal blood flow of both FLS and SLS groups**.

**S3-Full-field electroretinogram data sets. Excel file containing all data underlying the findings related to full-field electroretinogram of both FLS and SLS groups**.

